# Cross-generational characterization of paternal Low Protein Diet effects in mice

**DOI:** 10.64898/2026.10.16.759665

**Authors:** Flavio Santilli, Marco Riposati, Laura Castaldi, Bernhard Drotleff, Ana Boskovic

## Abstract

The impact of diet on the physiology of organisms and their progeny has been well documented across phyla. Using the established paternal Low Protein (LP) diet paradigm in mice, here we assess the robustness of the diet-induced intergenerational responses upon variation of genetic background and dietary history of tested animals. Protein restriction in FVB/NCrl resulted in mild, but specific hepatic responses in exposed animals, as well as their F1 progeny, affecting fatty acid metabolism and cholesterol biosynthesis, respectively. These phenotypes were qualitatively conserved between FVB/NCrl and C56BL/6J mice, suggesting the existence of shared molecular mechanisms underlying metabolic reprogramming, modulated in magnitude by genetic background. Unexpectedly, we uncover cumulative intergenerational inheritance effects that get reshaped at every generation, and a sex-specific ‘priming’ of cholesterol metabolism of LP-sired F1 offspring, evident upon dietary challenge. Together, our data, spanning multigenerational physiology, transcriptomics and lipidomics, indicate that the phenotypic consequences of paternal LP consumption depend on the duration and/or on the interaction of dietary exposure across generations, leading to distinct trajectories in hepatic energy metabolism.

## Introduction

The phenotype of a living organism is determined by the complex interactions between its genotype and the environments it experiences (GxE)^1–4^. Studies across phyla have established that variation in dosage and duration of biological (pathogen exposure, microbiome), physical (toxins, temperature, diet) and social (hierarchy, maternal care, socioeconomic status) environmental cues, impacts organismal health and disease risk^5,6^. Moreover, studies over the last two decades have provided evidence connecting preconception metabolic and physiological state of parents and offspring fitness and disease propensity ^7–9^ indicating that organismal traits are shaped not just by its own environment, but by its ancestors’ as well^10–13^.

Since the early laboratory studies into environmental contribution to biological inheritance, the use of inbred individuals has highlighted epigenetic mechanisms - including DNA methylation, histone modifications and long/small RNAs - as attractive candidates in mediating this phenomenon (reviewed in ^14,15^). Because these marks are dynamic and reversible, they allow organisms to adjust phenotypes to environmental conditions without changing the DNA sequence, thereby sitting at the interface of genome and environment to shape complex traits. Critically, epigenetic information can be transmitted through gametes, allowing environmental experiences to influence phenotypes across generations, a process known as epigenetic inheritance ^16,17^. Linking parental and grandparental preconception environments to descendants’ disease risk carries major evolutionary and biomedical implications, potentially explaining variation in disease onset and progression. In principle, this type of information transfer would allow the parents to ‘prime’ their progeny about the environments they might face, producing adaptive or maladaptive responses when expectations match or don’t match reality, as proposed by Gluckman and Hanson ^18,19^, and by Hales and Barker’s thrifty phenotype hypothesis^20,21^.

Given the universal nature of nutrition, the direct relevance to humans, and the ease of administration, dietary challenges (from starvation and/or caloric-restriction ^22^ to obesogenic and high-fat diet consumption^23,24^) have proven popular in the studies of epigenetic inheritance, in species ranging from worms^25,26^ to humans ^27,28^. One of the best characterized dietary paradigms in mammals is paternal Low Protein diet (referred to as LP henceforth): consisting of ad libitum consumption, from weaning to sexual maturity, of an isocaloric diet with reduced protein (∼10% instead of ∼20%) and increased carbohydrate (∼63% to ∼72%) content. Key studies have shown that chronic exposure of isogenic C57BL/6J male mice to LP predisposes their progeny to multi-organ metabolic alterations, including dysregulation of cholesterol biosynthesis in the liver^29,30^ and decreased lipogenesis in white adipose tissue^31^, observable already at early postnatal stages. These early-life alterations have been proposed to contribute to male-specific mild cardiovascular and metabolic complications in adulthood, such as reduced blood pressure, elevated heart rate, and impaired glucose clearance ^32^, with certain tissue-specific lipid responses reportedly persisting until F2^33^.

However, while the use of isogenic animals has been invaluable in dissecting purely environmental influence on phenotype across generations, it does not capture the complexity of biological inheritance, which is strongly influenced by genetic variation.

Our previous work uncovered an important role of genetic background in mediating immediate transcriptional responses in F1 blastocysts derived from sperm of males on either CHOW or LP ^34^. We observed notable differences in the baseline intrinsic gene expression variance, or ‘noise’, associated with C57BL/6J and FVB/NCrl mouse strains, pointing towards a strong interaction of genetics with environmental cues resulting in strain-specific transcriptional patterns as early as embryonic day 4. The FVB/NCrl genome carries approximately 4.3 million single-nucleotide polymorphisms (SNPs), and about 30,000 structural variants compared to C57BL/6J, including changes that affect the coding sequence of more than 400 genes^35^. Several of these occur within loci related to lipid metabolism, folate and one-carbon pathways, and cardiovascular function, suggesting that this strain’s metabolic wiring and responsiveness to dietary cues may differ markedly from that of C57BL/6J^35^, traditionally used in metabolic studies. In addition, FVB/NCrl is homozygous for Pde6b rd1, rendering the animals effectively blind, which might affect feeding behavior^36^. These observations prompted a direct investigation into the conservation of inheritance of adult phenotypes when studying GxE effects. Finally, it is still unclear whether paternal ‘priming’ of offspring metabolism confers adaptive advantage to the next generation, faced with the same or mismatched nutritional environment.

To systematically dissect the GxE effects related to paternal LP exposure, we established this paradigm in the FVB/NCrl mouse line following the original design principles, and asked the following questions: (i) how do FVB/NCrl males respond to dietary protein restriction?, (ii) are F1 metabolic phenotypes conserved between strains?, (iii) are F0 and F1 responses sensitive to ancestral dietary history, and ultimately (iv) what are, if any, the physiological consequences of paternal ‘priming’ on offspring, upon dietary challenge?

## Results

In order to test the impact of genetic variation on intergenerational inheritance induced by protein restriction, we set up the paternal LP paradigm in the inbred FVB/NCrl strain. Similarly to the previously established paradigm in C57BL/6J, two generations (2-gen) of breeding pairs were fed Control diet (henceforth CTRL) to washout CHOW food contribution, before exposing F0 sibling males to either CTRL or LP for 7-12 weeks, starting from weaning (Fig. 1A). The nearly-identical weight-gain trajectories and liver/body weight ratios between CTRL and LP-fed males was consistent with isocaloric dietary conditions (Fig. 1B, Fig. S1A, Supplemental Table 1). After dietary exposure, males from both groups were mated with sibling CTRL females, and subsequently sacrificed for liver transcriptomic and lipidomic profiling. Pregnancies were carried to term, and pups were weighed every 3 days from postanatal day 1 (P1) to weaning (P21), when they were euthanized for the same liver analyses.

**Figure. 1.**
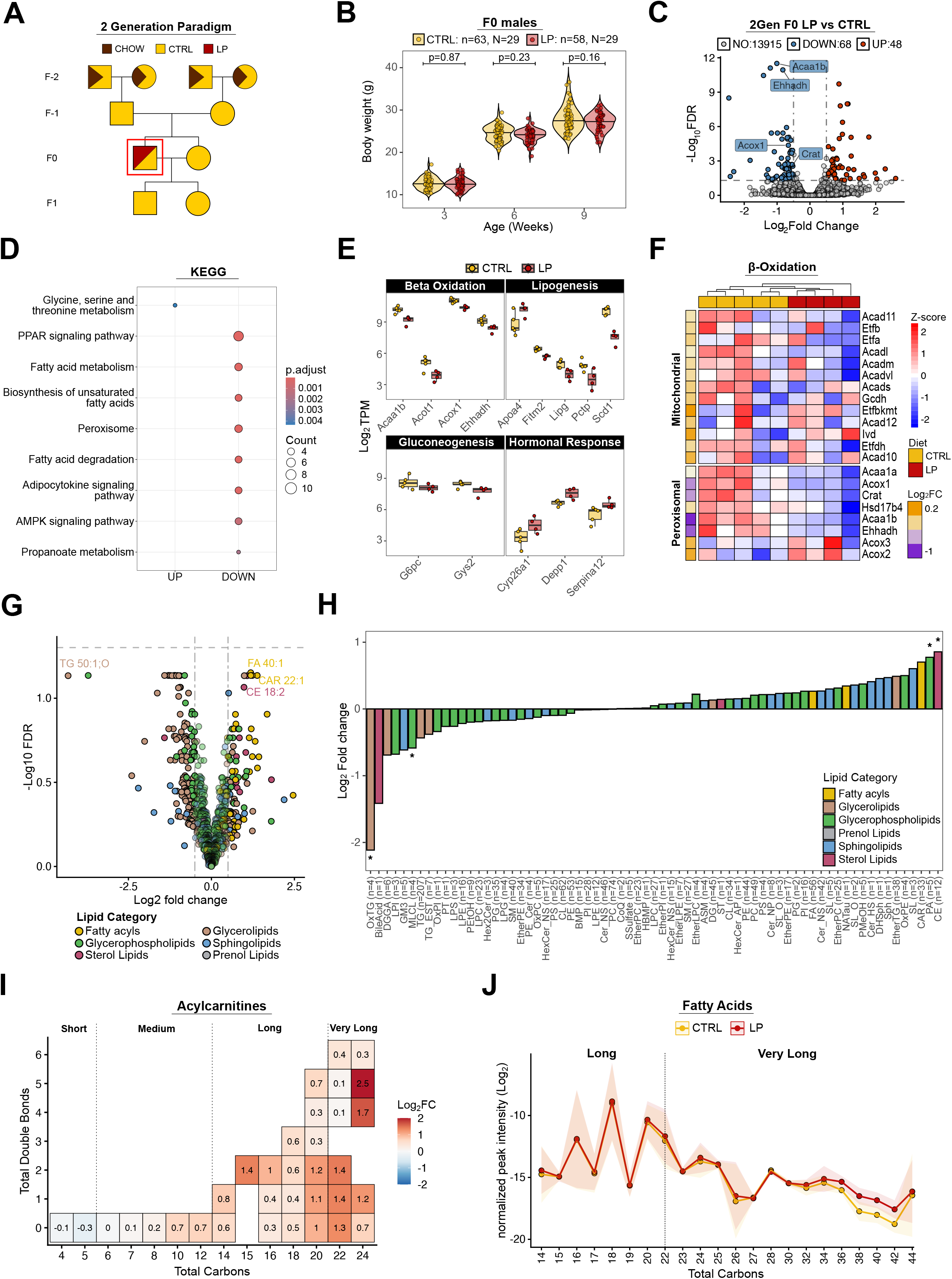
Impact of Low Protein Diet consumption on liver metabolism of FVB/NCrl mice. **A.** Breeding scheme and dietary intervention behind the 2-generation paternal LP paradigm. The box highlights the F0 generation analyzed. Squares and circles represent males and females, respectively. A bicolor triangle indicates a dietary transition (CHOW to CTRL), a diagonal split indicates the two alternative experimental groups (CTRL vs. LP). **B.** Violin plot of body weight measurements of F0 males fed CTRL diet or LP, from weaning for 6 weeks. Differences between experimental groups were assessed by a post-hoc analysis on a linear mixed model, considering animals in the same cage as non-independent (see Methods). **C.** Volcano plot of all detected genes from liver bulk RNA-seq data of F0 males, after filtering. Genes with adjusted p-values <0.05 & log2 Fold Change > 0.5, were considered significantly differentially expressed (DEGs) in LP vs CTRL testing. **D.** Dot plot of significant KEGG pathways enriched in DEGs, over all the detected genes. **E.** Boxplot of log2 Transcripts Per Million (TPM) of selected DEGs categorized into reported metabolic functions and grouped by diet. **F.** Heatmap of z-scores of selected genes involved in the mitochondrial or perixosomal β-oxidation. **G** Volcano plot of differential lipid abundance in LP vs. CTRL livers obtained through untargeted lipidomics. Each point represents an individual lipid species, colored by LIPID MAPS category. Dashed lines indicate log2 fold change (±0.5) and adjusted p-value (0.05) thresholds. **H.** Bar plot of class level log2 fold changes in lipid abundance in LP vs. CTRL samples. Different lipid categories are color-coded. Significant lipid classes are annotated (* adjusted p-value < 0.05). **I.** Heatmap of log2 fold changes of acylcarnitines grouped by total number of carbons and double bonds, highlighting compositional trends in the LP group. **J.** Line graph showing the mean (solid line) ± SD (shaded area) of normalized fatty acid intensities, grouped by number of carbon atoms.

We performed RNA sequencing of the livers from five pairs of male littermates fed CTRL or LP to chart the LP-effects on hepatic physiology in the parental generation. Principal component analysis (PCA) and linear regression modeling of the resulting data captured litter identity as the primary source of variance, with dietary conditions exerting a smaller but detectable influence on the transcriptomic variability (Fig.S1B-D). Differential gene expression analysis using DESeq2^37^ identified 116 differentially expressed genes (DEGs) (adjusted p-value <0.05, log2Fold Change > 0.5) between the two conditions (Fig. 1C, Supplemental Table 2). Among the downregulated genes, over-representation analysis on KEGG categories revealed an enrichment of transcripts encoding for factors involved in fatty acid metabolism (Fig. 1D, Fig. S1E). Curiously, genes involved in both *de novo* lipogenesis, lipid storage and fatty acid degradation were expressed at lower levels in LP livers (Fig. 1E). Since several β-oxidation factors (*Acox1, Ehhadh* and *Acaa1b*) were among the most significant DEGs, we next examined the expression of genes involved in either mitochondrial or peroxisomal β-oxidation pathways, which use Acyl-CoA dehydrogenases or Acyl-CoA oxidases in the first step of the cycle, respectively^38^. Interestingly, genes encoding peroxisome-specific enzymes, but not their mitochondrial counterparts, were expressed at lower levels in LP livers (Fig. 1F), pointing towards an impact on very long chain and branched fatty acids, specifically.

To connect our gene expression data to metabolic phenotypes, we next performed lipidomic profiling of the same liver samples. In agreement with the transcriptomic data, hepatic lipid composition largely overlapped between CTRL and LP fed males, with no individual lipid species showing significant changes (Fig. 1G, Supplemental Table 3). Nonetheless, through Lipid Set Enrichment Analysis (LSEA)^39^, we identified a significant coordinated decrease of triglycerides, and an enrichment of steryl esters and very long chain fatty acyls (Fig. S1F). Grouping individual lipid species into functional categories based on their chemical structure confirmed a significant accumulation of cholesteryl esters (CE) and a broad depletion of glycerolipids, particularly of oxidized triglycerides (OxTG), in the livers of LP F0 males (Fig. 1H, Fig. S1G). These observations are consistent with the strong transcriptomic downregulation of *Scd1* (Fig. 1E), suggesting a slowdown of *de novo* lipogenesis^40,41^. Interestingly, we observed a modest but consistent enrichment of fatty acyls in LP livers, specifically driven by increased levels of long and very long chain acylcarnitines and fatty acids (Fig. 1I-J). The observed chain-length-dependent impact of LP on acylcartinites might represent a dose-response pattern to the halving on dietary protein content. The concordance of transcriptomic and lipidomic data points towards slower degradation and/or decreased export, and consequent accumulation, of very long-chain fatty acids. Together, -omics analyses revealed a mild impact of LP-consumption on liver metabolism, with males on LP exhibiting a slightly slower turnover of very long chain fatty acids processed in the peroxisome, partially recapitulating previous reports in C57BL/6J strain^29,32^.

We next asked whether paternal LP consumption impacts offspring liver metabolism of FVB/NCrl mice as previously demonstrated in C57BL/6J ^29,30^ (Fig. 2A). Sex ratio (Fig. S2A) and average litter size (Fig. S2B) were not affected by paternal diet. Alongside, weight-gain across all tested groups was comparable from birth to weaning (Fig. 2B,C), with a significant increase in the liver to body weight ratio in LP-sired females at weaning (Fig. S2C). Liver transcriptomes of 12 pups sired across 3 litters obtained by pairs of sibling males on either CTRL or LP (N=3 litters, n=12 mice per experimental group) revealed that the largest source of transcriptomic variance (PC1) corresponded to cage identity, while PC2 captured sex differences, consistent with the reported strong liver sexual dimorphism (Fig. S2D,S2E). Paternal dietary status exerted modest but measurable impact on F1 hepatic gene expression, and we identified 130 DEGs when comparing LP-sired with CTRL-sired F1 livers (Fig. 2D, Supplemental table 2). Gene Ontology (GO) Term analysis of downregulated genes did not highlight any particular functional pathway. In contrast, among upregulated genes, we found significant enrichment for multiple terms related cholesterol biosynthesis (Fig. 2E), including 9/23 core biosynthetic enzymes (*Acat2, Dhcr7, Nsdhl, Mvk, Idi1, Tm7sf2, Sqle, Cyp51, Sc5d*) (Fig. 2F) ^42^. Altogether, strong upregulation of *Acly*, supporting acetyl-CoA production, and of the two rate limiting steps (*Sqle, Hmgcr)* in both males and females F1 (Fig. S2F), pointed to enhanced *de novo* cholesterol synthesis in the livers of LP-sired mice. Curiously, *Srebf2,* encoding sterol regulatory element-binding protein 2 (SREBP-2), normally activated by low cholesterol levels and positively regulating *Hmgcr* transcription, was not differentially expressed. Intersecting our transcriptomic data with publicly available dataset from C57BL/6J mice on the same protein restriction paradigm (45% protein reduction compared to CTRL) ^29^ revealed a strong overlap between the upregulated genes across the two strains (Fig. 2G). Enrichment analysis on overlapping genes (Fig. S2G) confirmed that the functional metabolic consequences of paternal protein deficiency are conserved between strains, laboratories and transcriptome detection methods (microarray versus RNA-sequencing), reinforcing the robustness of intergenerational metabolic reprogramming in response to paternal dietary conditions in mammals.

**Figure. 2.**
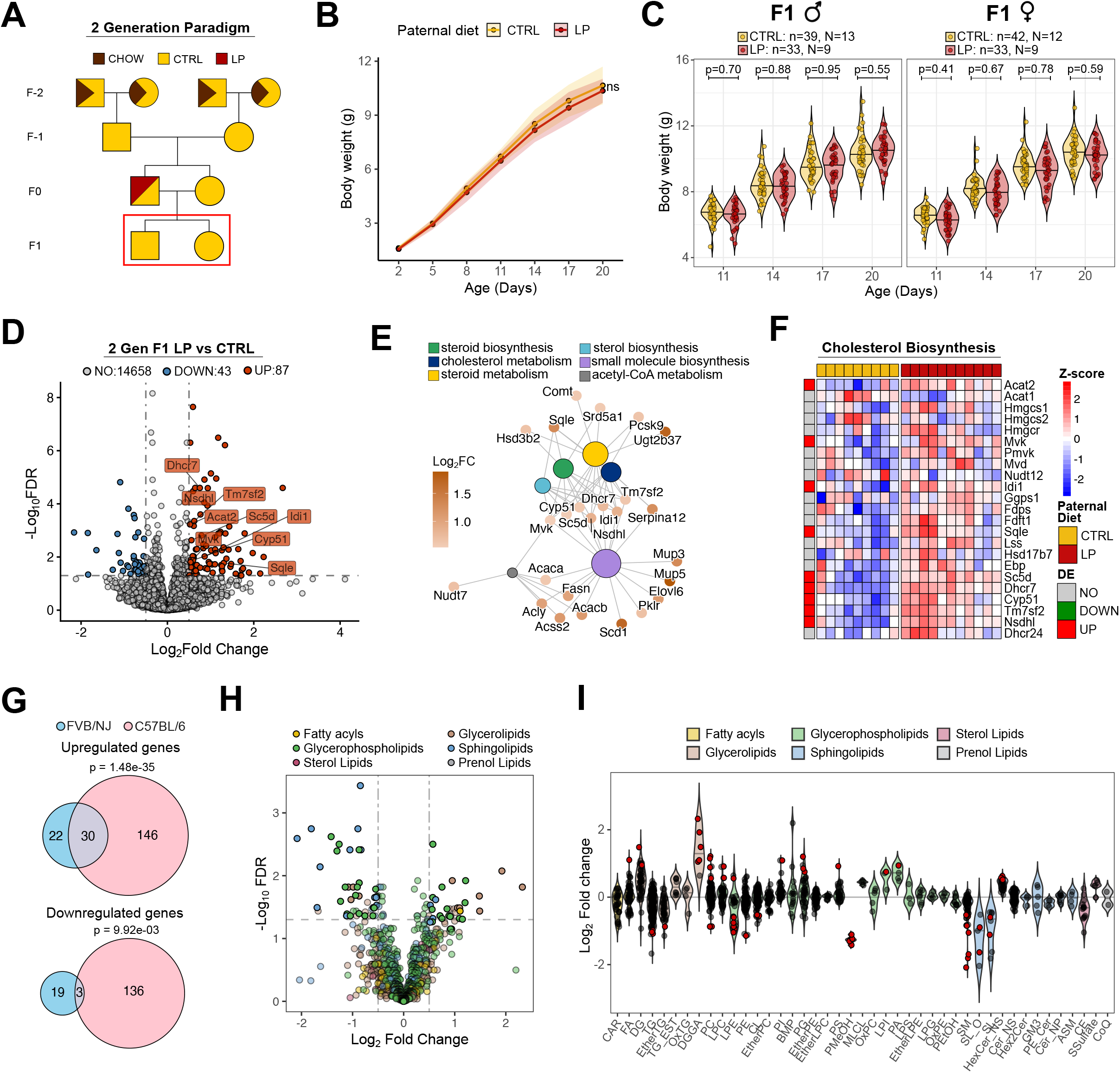
Intergenerational impact of paternal LP consumption in FVB/NCrl. **A.** Breeding scheme and dietary intervention behind the 2-generation paternal LP paradigm. The box highlights the F1 generation analyzed. Squares and circles represent males and females, respectively. A bicolor triangle indicates a dietary transition (CHOW to CTRL), a diagonal split indicates the two alternative experimental groups (CTRL vs. LP). **B.** Growth curves of pups sired by CTRL and LP-males, from birth to weaning. Difference in growth dynamics between experimental groups was assessed by a post-hoc analysis on a linear mixed model, considering animals in the same cage as non-independent (see Methods). **C.** Violin plot of body weight measurements of pups sired by CTRL and LP males, from postnatal day 11 to 20, stratified by sex. Differences between experimental groups were assessed by a post-hoc analysis on a linear mixed model, considering animals in the same cage as non-independent (see Methods). **D.** Volcano plot of all detected genes from liver bulk RNA-seq data of F1 pups, after filtering. Genes with adjusted p-values <0.05 & log2 Fold Change > 0.5, were considered significantly differentially expressed (DEGs). **E.** Gene concept network of the 6 most significant GO terms identified by over-representation analysis of DEGs, with genes colored by log2 Fold Change. **F.** Heatmap of Z-scores for cholesterol biosynthesis genes. Experimental groups and statistically significant features are color-coded. **G.** Venn diagrams of F1 Upregulated (top) and Downregulated (bottom) DEGs from this study (FVB/NCrl) and Carone 2010 [34] (C57BL/6J). **H.** Volcano plot of differential lipid abundance in F1 LP vs. CTRL livers obtained through untargeted lipidomics. Each point represents an individual lipid species, colored by LIPID MAPS category. Dashed lines indicate log2 fold change (±0.5) and adjusted p-value (0.05) thresholds. **I.** Violin plot of Log2 Fold Changes of single lipids grouped by LIPID MAPS ontologies. Red points highlight significantly differential lipids.

Lipidomic analyses of the same samples revealed that only ∼6% (73/1332) of detected lipids differed statistically between LP and CTRL F1, with the majority of them showing only a modest change (1 < log2 fold change > -1) (Fig. 2H, Supplemental Table 3). Many significant enriched lipids were diglycerides (DGs) (Fig 2I, Fig. S2H), in line with the intergenerational effects reported in C57BL/6J, and possibly supported by the upregulation of key lipogenic genes (*Fasn, Scd1, Acly*). Conversely, approximately half of lysophosphatidylethanolamines (LPE) were depleted in F1 LP-livers. Surprisingly, overall levels of cholesteryl esters (CE) were comparable between F1 LP and CTRL groups, in contrast to the results from C57BL/6J and the transcriptomic signatures observed in our data (Fig. 2I), suggesting that increased transcriptional flux through the pathway may not be reflected in steady-state cholesterol pool. This discrepancy could reflect increased cholesterol turnover or export (e.g., to plasma) that offsets synthesis, or may represent a transient transcriptional response that has already been compensated for by weaning. Notably, lipid changes reflected some degree of differential response from the three different litters analyzed (Fig. S2I), pointing towards the partial penetrance of intergenerational LP phenotypes in FVB/NClr.

Environmental conditions impacting mammalian physiology are dynamic in nature, and typically do not persist over generational time. Thus, to understand the sensitivity of LP-mediated metabolic reprogramming to environmental shifts, we shortened the duration of ancestral CHOW to CTRL diet wash-out time from two (F-2, or 2-gen) to one (F-1, or 1-gen) generation, before exposing the experimental F0 males to either CTRL or LP consumption (Fig. 3A).

**Figure. 3.**
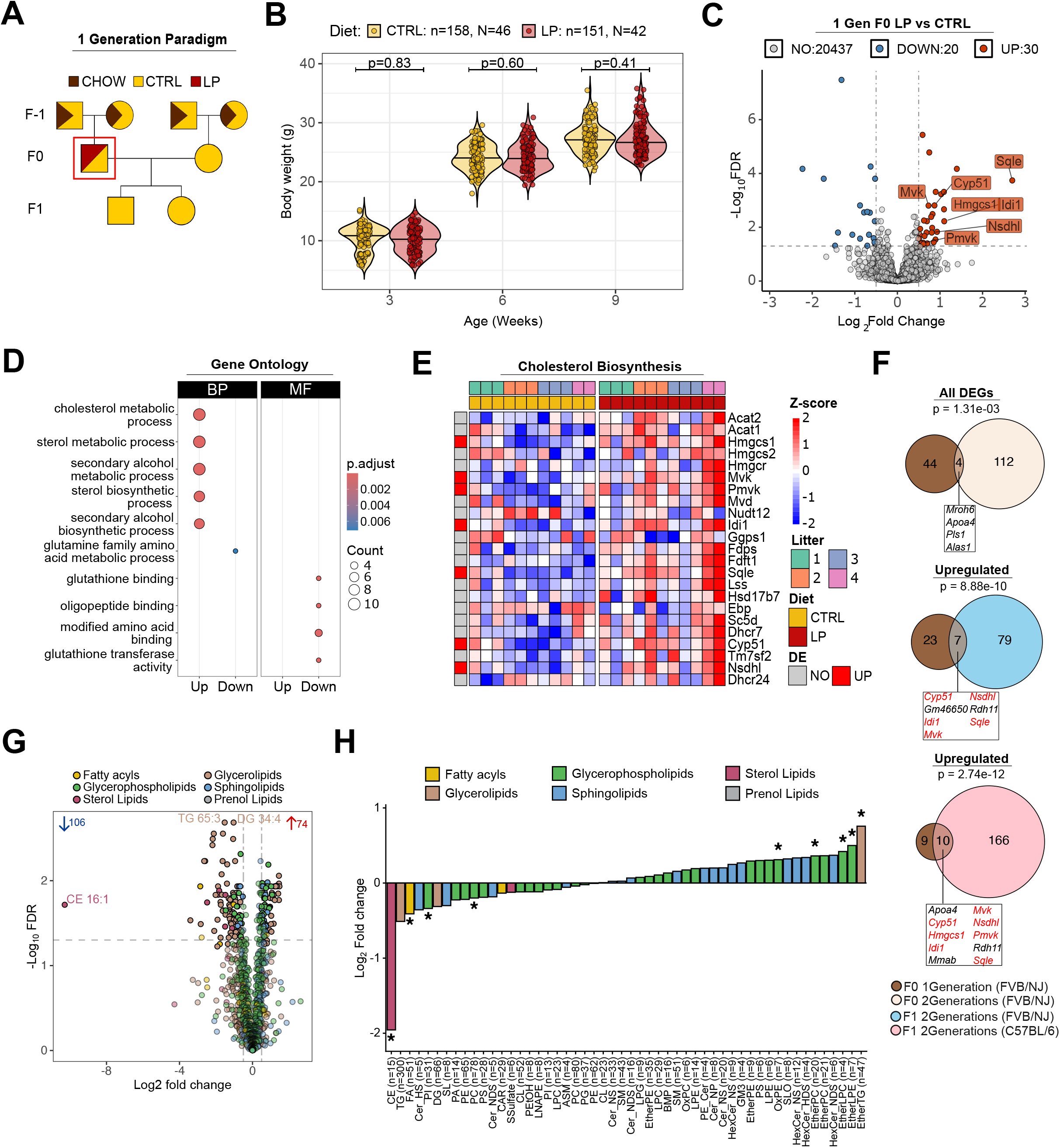
Liver profiling of LP exposed animals in the 1-generation Paradigm. **A.** Schematic of 1-generation (1-gen) Control washout Low Protein diet paradigm. The box highlights the F0 generation analyzed. Squares and circles represent males and females, respectively. A bicolor triangle indicates a dietary transition (CHOW to CTRL), a diagonal split indicates the two alternative experimental groups (CTRL vs. LP). **B.** Violin plots showing single animal weights at indicated time points. Animal (n) and cage (N) numbers are indicated. Differences between experimental groups within individual time points were tested using a linear mixed model controlling for multiple covariates (see Methods). **C.** Volcano plot showing the result of differential gene expression analysis of LP vs CTRL F0 livers. Number of DEGs (adjusted p-value < 0.05 & absolute log2 fold change > 0.5) are indicated. Upregulated genes belonging to the cholesterol biosynthesis pathway are labeled. **D.** Dotplot of top 10 Gene Ontology (GO) terms enriched in up- and downregulated DEGs. BP = biological processes, MF = molecular functions. **E.** Heatmap of cholesterol biosynthesis pathway genes, showing scaled expression (z-scores) across biological replicates under CTRL and LP regimes. **F.** Venn diagrams of DEGs comparing the effects of LP across different dataset. **G.** Volcano plot of LP vs CTRL hepatic lipids colored by category. Significant lipids (adjusted p-values <0.05 & log2 Fold Change > 0.5) are annotated. **H.** Average log2 Fold change (LP vs CTRL) across lipids subclasses colored by Lipid MAPS category. Statistical significance was assessed using a moderated t-test with Benjamini–Hochberg correction (FDR) (* = adjusted p-value < 0.05).

Consistent with the findings of the 2-gen paradigm (Fig. 1B), consumption of LP did not impair the weight gain of males at the three measured timepoints (Fig 3B). PCA on liver transcriptomic data highlighted a cage-based clustering together with some degree of separation driven by the different diets consumed (Fig. S3A). Statistical testing of males fed CTRL or LP (N=4 litters, n=12 animals per condition) revealed that LP consumption induced weak general transcriptional changes, with only 50 genes reaching statistical significance of differential expression (Fig. 3C, Supplemental Table 2). Strikingly, alongside a mild association of downregulated genes with protein clearance, we detected a significant enrichment of genes responsible for cholesterol biosynthesis among the upregulated DEGs (Fig. 3D, Fig. S3B, C). In addition, genes belonging to this metabolic process, whose expression difference did not reach statistical significance, showed a trend of increased expression in LP-fed males, pointing towards a modest, but consistent enhancement of cholesterogenesis (Fig. 3E). Surprisingly, comparing DEGs detected in F0 livers from 1-gen and 2-gen animals revealed only four commonly deregulated genes (p = 0.001). Conversely, a more significant overlap was observed when comparing F0 1-gen LP-induced changes to those of F1 2-gen samples, from both FVB/NCrl (p =8.88e-10) and C57BL/6J (p = 2.7e-12) genetic background (Fig. 3F). This overlap, enriched for cholesterol biosynthetic transcripts, occurred specifically between upregulated, but not downregulated genes (Fig. 3F, Fig. S3D), highlighting enhanced cholesterogenesis as the only shared transcriptional signature.

Unlike the observations in the 2-gen paradigm, lipidomic analysis of 1-gen liver samples revealed substantial differences in lipid content between dietary groups, significantly affecting 180 lipids (Fig. 3G, Supplemental Table 3). Even more curious was the striking depletion of cholesteryl esters (CE) in LP-livers (Fig. 3H), possibly underlining a state of negative hepatic cholesterol balance triggering the concomitant upregulation of pivotal genes involved in *de novo* cholesterol biosynthesis such as *Srebf2*, *Hmgcs1*, *Sqle* (Fig. 3E, Fig. S3B)^43^. Noteworthy, this finding in F0 1-gen males mirrors the phenotype previously reported in the offspring (F1) of LP fed fathers in C57BL/6J strain^29^. In parallel, fatty acids and their esterified glycerolipids (di- and triglycerides) were consistently depleted, particularly very-long-chain and unsaturated species (Fig. S3E), suggesting a shift of lipid allocation given the generalized increase of ether lipids (Fig. 3H). Of note, the magnitude of transcriptional (Fig. S3B) and lipidomic (Fig. S3F) responses to LP was litter-dependent, ranging from mild (Litter 1) to pronounced (Litter 2).

Collectively, these results corroborate that consumption of LP from weaning to sexual maturity modestly impacts liver metabolism of male mice, but highlight how ancestral dietary history plays an important role in determining specific responses to metabolic shift in mammals, hinting towards a multigenerational interaction of dietary information in determining metabolic traits.

Given the enhanced cholesterogenesis in F0 males (1-gen) (Fig. 3D, S3C) and F1 offspring of LP-fed sires (2-gen) (Fig. 2C), we asked whether this paternal LP-induced upregulation of cholesterol biosynthesis persists across generations after a single generation of control diet wash-out (Fig. 4A). The pre-weaning growth dynamics of F1 pups from CTRL and LP sires was comparable, across both males and females (Fig. 4B, Fig. S4A), as well as their sex ratio and litter size distribution (Fig. S4B,C). Transcriptomic analysis of offspring liver samples (N=4 litters, n=16 animals, for either paternal diet) revealed that sex differences are the main source of transcriptomic variation in the data, as before (Fig. S4D, Supplemental Table 2). To understand the influence of paternal diet on hepatic mRNA profiles, we visualized the sample distribution along the principal component axes (Fig. S4D,E). We noted a mild effect of paternal LP exposure, and identified 61 downregulated and 73 upregulated genes through differential gene expression analysis (Fig. 4C). Notably, key regulators of phase I (*Cyp3a41a, Cyp3a44, Cyp4a31*) and phase II detoxification processes (*Ugt1a7c, Ugt1a8, Sult3a1*) showed coordinated increase in gene expression (Fig. 4D, E)^44,45^. Downregulation of *Fasn* (Fatty acid synthase) and *Acly* (ATP citrate lyase), and *Scd1* (Stearoyl-CoA Desaturase 1) suggested a slower rate of *de novo* fatty acid synthesis in LP-sired offspring^46,47^. Together with the increased levels of *G6pc* encoding glucose-6-phosphatase, a pivotal enzyme in glucose homeostasis, these data point to an early metabolic reprogramming that favors carbohydrate utilization over lipid synthesis (Fig. 4E). Of note, despite this metabolic rewiring, in the 1-gen paradigm paternal LP did not significantly impact any of the cholesterol biosynthesis transcripts (Fig. S4F), highlighting a differential influence of paternal LP on de novo lipogenesis and cholesterogenesis across generations, paradigms and strains. To validate this, we calculated the Spearman correlation of the relative expression changes (log2FC) among experimental groups for cholesterol (left) and general lipid biosynthesis (right) gene sets (Fig. 4F). For the former, F0 1-gen LP-driven transcriptomic changes correlated strongly with both F1 2-gen FVB/NCrl (r=0.66, p=<5.89 e-05, 95% CI [0.41, 0.81]) and published F1 C57BL/6J data (r=0.60, p=0.01, 95% CI [0.12, 0.90]), while correlation with F0 2-gen was weaker (r=0.44, p=0.01, 95% CI [0.11, 0.68]). The same pattern was observed for general lipid biosynthesis, though of lower magnitude (F0 1Gen∼F1 2Gen FVB/NCrl: r=0.18, p=2.67e-05; F0 1Gen∼F1 2Gen C57BL/6J: r=0.37, p=5.26e-04, F0 1Gen∼F0 2Gen: r=0.01, p=0.73). In contrast, conservation of intergenerational responses across different genetic backgrounds (within the 2-gen paradigm context) was widespread, with high correlations in both cholesterol (r = 0.61, p = 0.009) and general lipid biosynthesis (r = 0.76, p = 6.49e-17). Even more interestingly, transcriptional outputs in F1 1-gen showed unique features compared to all the other permutations of the paradigm, generation and strain, indicating a reshaping of metabolic responses with each successive dietary change. This pattern was further supported by the groups identified through unsupervised clustering of gene-expression changes for individual cholesterol biosynthesis genes across generations and genetic backgrounds (Fig. 4G).

**Figure. 4.**
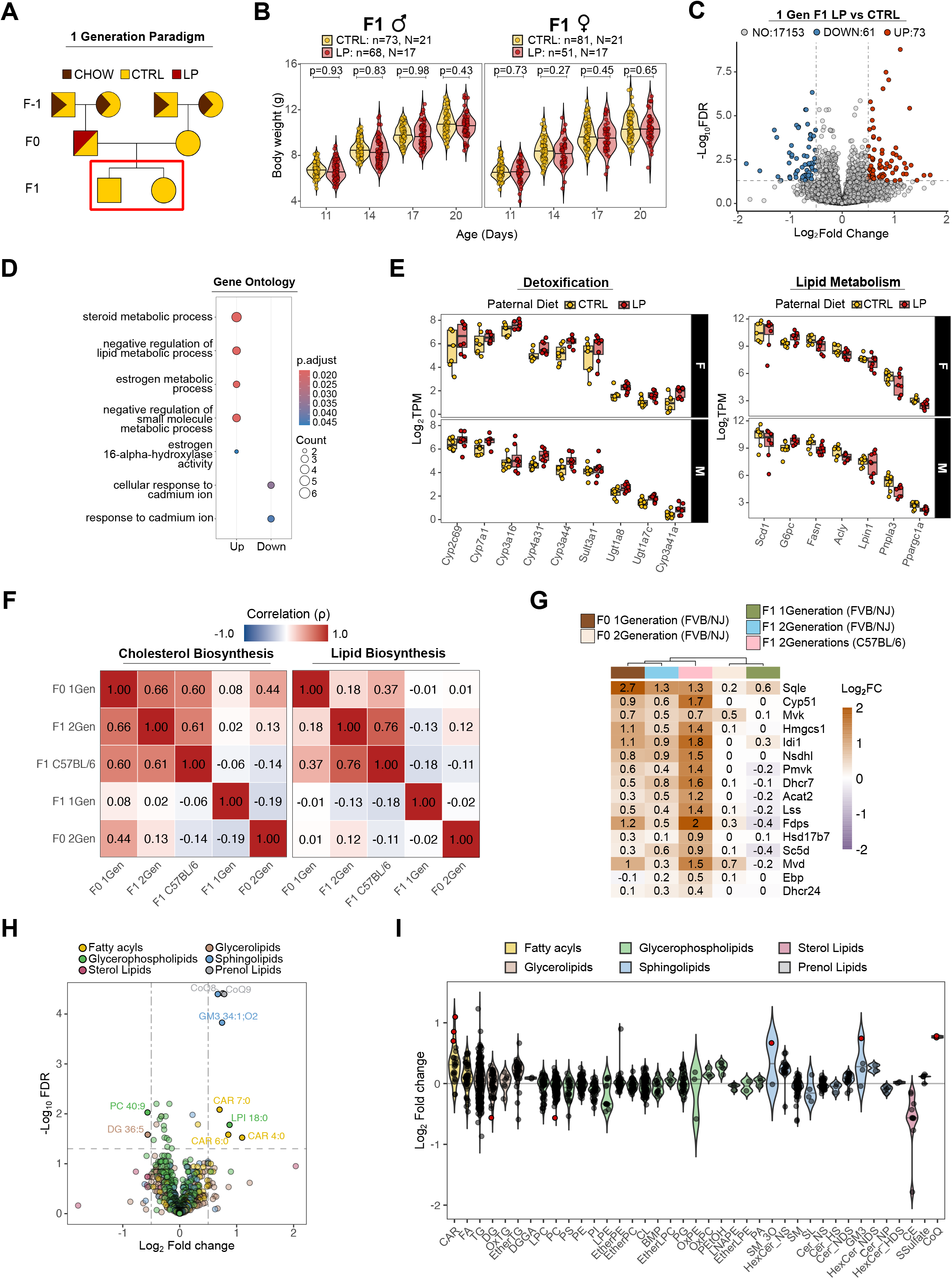
Intergenerational effects of paternal LP in the 1-generation paradigm. **A.** Schematic of 1-generation (1-gen) Control washout Low Protein diet paradigm. The box highlights the F1 generation analyzed. Squares and circles represent males and females, respectively. A bicolor triangle indicates a dietary transition (CHOW to CTRL), a diagonal split indicates the two alternative experimental groups (CTRL vs. LP). **B.** Violin plots of body weights from single male and female offspring at indicated postnatal stages. Animal (n) and cage (N) numbers are shown. Statistical differences between groups were assessed using t-tests on a fitted linear mixed-effects model accounting for litter as a random effect (see Methods). **C.** Volcano plot of differential expression analysis of LP vs CTRL F1 livers. The number of DEGs (adjusted p-value < 0.05 & absolute log2 fold change > 0.5) are indicated. **D.** Dotplot of significantly enriched Gene Ontology functional terms among F1 LP vs CTRL DEGs. **E.** Boxplot of normalized expression levels (log2 TPM) of representative detoxification and lipid metabolism genes across individual samples grouped by biological sex and diet. **F.** Spearman correlation matrices comparing LP vs CTRL log2 FC across experiments for cholesterol biosynthesis (left) and general lipid (right) gene sets. **G.** Heatmap of log2 Fold changes of commonly detected cholesterol biosynthesis genes across the different dietary paradigms, generations, and genetic backgrounds. **H.** Volcano plot showing differentially abundant lipids (adjusted p-value < 0.05 & absolute log2 fold change > 0.5) between F1 LP vs CTRL livers. **I.** Violin plot of log2 FC of single lipids, grouped by subclasses. Red points represent significant differential lipids.

Examining F1 1-gen lipid profiles from different paternal dietary groups revealed that they largely overlapped, with only 10 lipid species reaching statistical significance of differential abundance when comparing LP and CTRL-sired offspring (Fig. 4H, Supplemental Table 3). Grouping individual lipids by LIPID MAPS^48^ classes revealed a mild increase of shorter chain acylcarnitines (Fig. S4G), and a consistent enrichment of coenzyme Q (CoQ) in LP F1 livers, which aligns with the alteration of detoxification-related genes, given the antioxidant activity of reduced forms of CoQ (CoQH2) (Fig. 4I) ^49,50^. Curiously, even when stratifying by sex or collection date (Fig. S4H), we did not detect any enrichment of DGs in F1 offspring of LP-consuming sires, contrasting the 2-gen lipidomic results. The divergence of F1 hepatic responses to paternal LP, depending on their ancestral dietary history, strengthens the notion that the metabolic consequences of paternal dietary protein restriction depend on the duration and/or on the interaction of dietary exposure across generations, leading to distinct trajectories in hepatic energy metabolism.

Finally, we set out to test if paternal dietary conditions confer (mal)adaptive properties to F1 liver metabolism. At weaning, F1 pups from CTRL or LP-fed sires on 2-gen paradigm were fed one of the three diets: CTRL, LP or High-Cholesterol (HCHOL) (see Methods and Supplemental Table 1), *ad libitum*, for 7-8 weeks (Fig 5A). These dietary conditions represented neutral, matched and mismatched environments, in relation to those experienced by the sires. Measurements of animal growth did not reveal any significant effect of paternal diet on body weight at individual time points, although we noted CTRL-sired mice tended to be slightly heavier, exhibiting significantly faster growth trajectories in males (Fig. S5A). Interestingly, F1s on HCHOL exhibited paler livers (Fig. S5B) coupled with increased liver/bodyweight ratio compared to the other two exposure groups (Fig. S5C), indicating a strong influence of dietary cholesterol supplementation on liver function. Although paternal diet did not strongly affect liver weight when comparing within the three different dietary challenges (Fig. 5B), LP-fed mice displayed significantly lower liver/body weight ratio compared to CTRL-fed F1, when sired by LP males (Fig. S5C). These results point towards an interaction of paternal and offspring dietary status in the context of protein restriction. We next performed RNA-seq on livers from tested animals (N=3 litter, n=5-6 animals per experimental group) to assess the interaction between paternal and consumed diets. PCA of the resulting data indicated that biological sex and consumed diet drive the sample separation along the first two principal components (Fig. 5C), highlighting the dominant effects of consumed diet over paternal diet in determining transcriptomic responses. Indeed, through differential expression analysis, we identified hundreds of significant DEGs when comparing the effects of F1 diets (Fig. S5D, Supplemental Table 2), but virtually no significant hits when assessing the transcriptional changes induced by paternal LP diet in any of the F1 dietary contexts (Fig. S5E). To capture coordinated contribution of paternal diet to offspring metabolism, we first performed k-means clustering of relative changes induced by paternal-consumed diet interaction. Next, we performed Over Representation Analysis (ORA) on the groups of filtered genes identified (absolute log2Fold Change > 0.5) (Fig. 5D), uncovering a bias towards cholesterol metabolism among the genes selectively activated by paternal LP, exclusively in the F1 mice eating an LP (cluster 3). Furthermore, when F1 mice sired by LP males were fed an HCHOL diet, *Mups* (associated with the odorant binding term) and NF-κB signaling pathway transcripts were enriched within the clusters of activated (cluster 5) and suppressed genes (cluster 6), respectively (Fig. 5D, E). Independently, Geneset Enrichment Analysis (GSEA) of paternal LP transcriptional effects confirmed F1 diet-specific signatures, but also revealed shared pathway alterations, including coordinated reduction of immune-related and activation of odorant binding genes, specifically when F1 mice were fed LP or HCHOL (Fig. 5F). Notably, these shared molecular signatures were consistent with the mild correlation between the paternal LP-induced transcriptional changes in LP- and HCHOL-fed offspring compared to those consuming CTRL diet, for which no correlation was detected (Fig. S5F). Given the significant activation of cholesterol metabolism in LP-sired LP-fed F1 and previous evidence of paternal LP in priming F1 cholesterol biosynthesis, we inspected the relative changes induced by paternal LP on individual genes involved in this pathway. Transcripts related to cholesterol biosynthesis (*Sqle, Idi1 Hmgcs1*), but not those involved in cholesterol sensing (*Srebf1, Srebf2*) or catabolism (*Cyp27a1, Cyp46a1*), were consistently induced by paternal LP, in both LP and CTRL consuming F1s (Fig. 5G). Intriguingly, stratifying the analysis by sex revealed a marked sexual dimorphism in transcriptional response to paternal LP. In both CTRL and LP F1s, females displayed a coordinated induction of cholesterol biosynthetic genes, while males showed comparatively limited, if not opposite changes (Fig. 5H). Together, these analyses reveal an unexpected interaction between paternal preconception and encountered dietary information in contributing to liver metabolism, additionally modulated by biological sex.

**Figure. 5.**
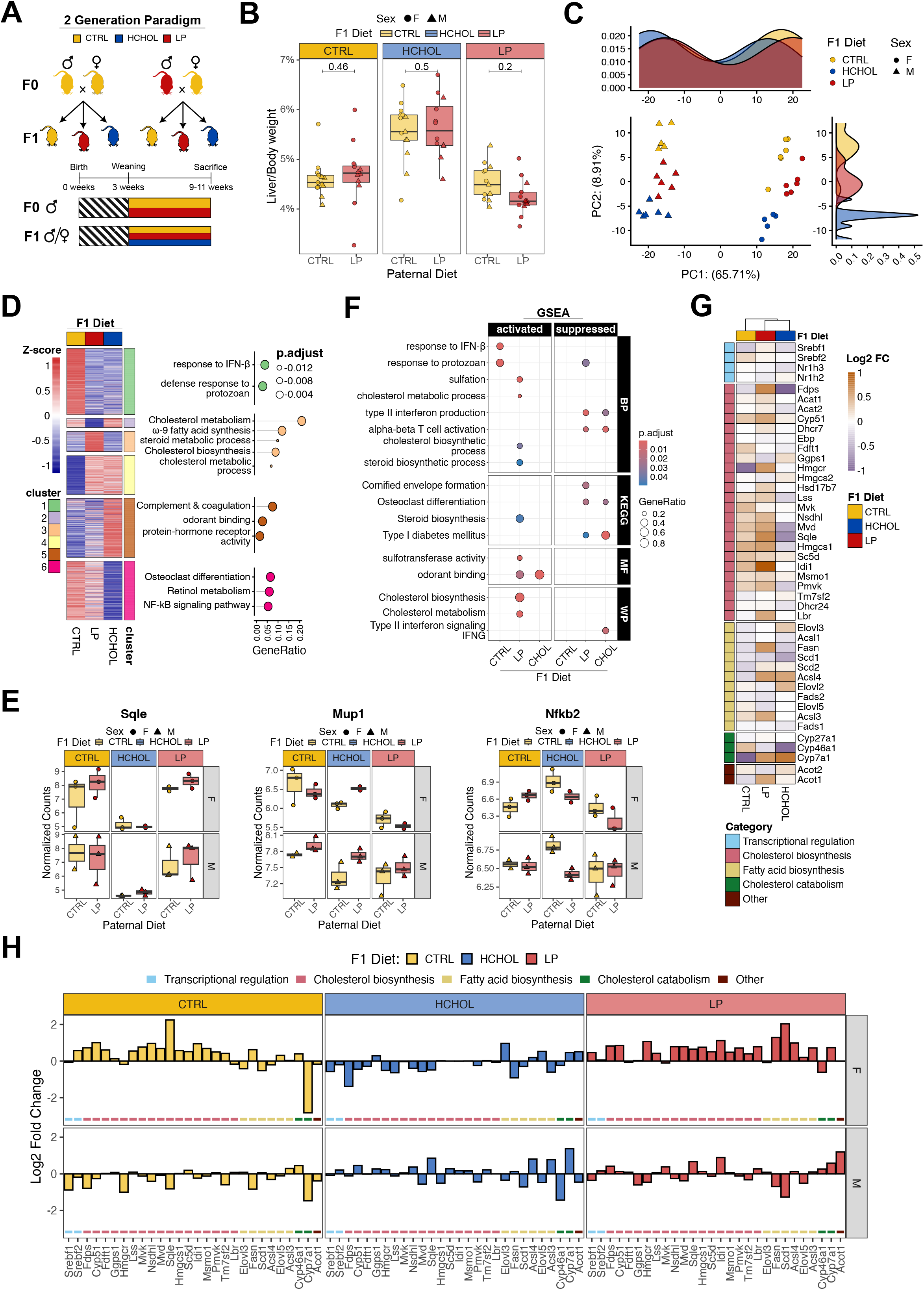
Impact of paternal diet on F1 hepatic signatures upon distinct dietary conditions. **A.** Schematic of the F0 and F1 dietary challenge experimental design. **B.** Boxplots of liver to body weight ratios of F1 animals derived from CTRL or LP-fed males, at the end of dietary challenge. Data are stratified by F1 consumed diets. The p-values were calculated using a two-tailed Wilcoxon rank-sum test. **C.** PCA of top 500 most variable genes in F1 liver RNA-seq samples shows separation based on sex (PC1) and diet consumed (PC2). **D.** Heatmap of gene Z-scores of log₂ fold changes calculated for each paternal LP effect across the three F1 contexts, grouped by k-means clusters. The dotplot on the right shows selected GO terms significantly enriched in the clusters indicated by the corresponding color fill. **E.** Boxplots of normalized expression levels (vst) of representative genes belonging to different clusters identified in D. Data was stratified by F1 diet consumed, and biological sex. **F.** Dotplot of significant terms identified by GSEA of paternal LP vs. CTRL comparisons stratified by F1 diet consumed. BP = biological processes. MF = molecular function, KEGG = Kyoto Encyclopedia of Genes and Genomes. WP = WikiPathways. **G.** Heatmap of log2 Fold changes of paternal LP across the F1 dietary contexts indicated at the top, showing genes involved in cholesterol metabolism color-coded by their specific subprocess. **H.** Barplot showing paternal LP-induced relative changes of selected genes involved in cholesterol metabolism, stratified by F1 diet and biological sex.

## Discussion

In this work, we systematically tested the impact of LP in FBV/NCrl mouse strain across generations, experimental paradigms and postnatal metabolic challenges. Our data revealed a conserved impact of paternal protein restriction on fatty acid (F0) and cholesterol (F1) metabolism between the FVB/NCrl and C57BL/6J strains, with an overall lower magnitude consistently observed in the FVB/NCrl animals. This is in good agreement with recent reports showing that despite higher baseline glucose levels, FVB mice develop glucose intolerance and increased adiposity under a high-fat diet, albeit to a different extent than C57BL/6J^51^. In addition, the dietary protein content used in our study (16.5% for CTRL and 9.1% for LP, 45% reduction) is not identical to that used previously (20% for CTRL and 11% for LP, 45% reduction), possibly contributing to subtle differences in phenotype strength. It is also noteworthy that the LP-consumption protocol modulates only the dietary content, but not the total caloric count or food availability. The subtlety of environmental challenge is likely reflected in the modest metabolic adjustments, rather than generalized reprogramming, we observed in our LP samples.

Although the coordinated transcriptional upregulation of *de novo* cholesterol biosynthesis is observed in F1 offspring of LP-fed males in both strains (FVB/NCrl and C57BL/6J), lipidomic analyses showed that diglycerides, but not cholesteryl esters, were predominantly affected in response to paternal LP in the FVB/NCrl background. The consistent increase in DGs, partially shared by C57BL/6J, may explain the upregulation of the lipogenesis transcriptional program, and has been associated with insulin resistance and Metabolic dysfunction-Associated Steatotic Liver Disease (MASLD)^52^. Whether these changes observed in pups at weaning contribute to metabolic impairments later in adulthood needs to be experimentally tested. Altogether, the findings from F0 and F1 generations demonstrate that FVB/NCrl mice recapitulate the transcriptional hallmarks of paternal LP reported in C57BL/6J, albeit with reduced magnitude. This, in turn, suggests the existence of a shared intergenerational mechanism of metabolic reprogramming, modulated in magnitude by genetic background.

An unanticipated finding emerging from this work is the profound impact of dietary history preceding LP consumption on metabolic phenotypes in mice. Reducing the number of generations undergoing a CTRL-diet washout from CHOW food from two to one resulted in markedly different metabolic responses in both F0 and F1 livers. These differences might arise from the interaction of dietary cues triggered by sequential shifts in the protein-to-carbohydrate ratio (from CHOW to CTRL, then from CTRL to LP), which act as nutritional signals that prime hepatic metabolism, and presumably the physiology of other organs. However, as the 1-gen and 2-gen cohorts were run separately, and the 1-gen F0 RNA-sequencing libraries were prepared externally, we note the possibility that the difference between transcriptomic outcomes may partly be confounded by batch effects. Since all lipidomic data were generated in a single facility, and also shows divergent profiles between the two cohorts, we speculate that the batch-effects constitute only a minor, if any, contribution to the measured molecular phenotypes.

Strikingly, when comparing the F1 offspring of LP-males with different ancestral histories, we observed very little overlap in their molecular signatures. It is plausible that cholesterol-related phenotypes already established in the fathers buffer or mask further intergenerational effects, pointing to a metabolic interaction between generations, in which the parental physiological state constrains the transmission of new dietary responses. By extension, males exhibiting strongest cholesterol biosynthesis responses upon LP-challenge would sire offspring with the least pronounced hepatic phenotypes, a hypothesis which requires testing across a large-scale experimental cohort considering the overall low magnitude of transcriptional and lipidomic responses to LP.

Finally, through our F1 dietary challenge experiment, we found that the diet consumed by the F1 animals outweighed the contribution of preconception paternal programming on liver transcriptomes. Nonetheless, we showed that paternal LP modulates the expression of metabolic and immune-related genes, depending on the offspring diet. This supports a model where the overall hepatic phenotype arises from the cross-generational interaction between inherited and experienced nutritional information. We speculate the additive intergenerational effects may be propagated further, as the F1 animals with different physiological states likely harbor specific epigenetic signals in their gametes, contributing to the phenotypes of their offspring. Further parallelized characterization of F1 metabolism, physiology, and gamete epigenomic signatures, coupled to targeted functional experiments, will be necessary to formally test this prediction. Finally, it is interesting that the transcriptional rewiring of cholesterol metabolism depended not only on the multigenerational dietary regimen, but also biological sex. LP-sired females showed a more prominent effect compared to their male siblings, although these findings require further validation across a larger experimental cohort. Indeed, sex-specific impact of parental programming has been previously reported in the literature^5,24,53–55^. This is an important consideration in experimental design, particularly relevant when assessing phenotypes in organs with strong molecular signatures of sexual dimorphism, such as the placenta, liver or brain.

Collectively, our work indicates that paternal LP-effects depend on the interaction between ancestral nutrition and metabolic state, determining whether metabolic reprogramming signals are transmitted, reshaped, or erased across generations. A critical open question remains: what are the underlying mechanisms? The overall low magnitude of differences, and phenotypic penetrance between offspring derived from males on distinct diets, is compatible with the notion of sperm-borne epigenetic factors acting as mediators of paternal effects, as has been proposed before^30,56^. However, additional experiments using assisted reproductive technologies (such as IVF) and F1 cross-fostering, as well as targeted functional perturbations, would be required to formally test this hypothesis. If indeed this is the case, it is likely that the ultimate adult phenotype arises from the combinatorial input of different information carriers and their interaction with oocyte (epi)genetics, and maternal physiology. Future mechanistic studies, powered by advances in precision molecular measurements in the germline, and strong statistical frameworks, will undoubtedly shed light on this outstanding question.

## Acknowledgments

We thank the members of the Boskovic group for insightful discussions and comments; the Laboratory Animal Resources at EMBL Rome, particularly Olga Boruc, for assistance with animal husbandry and procedures; the EMBL Genomics Core Facility, particularly Vladimir Benes, for support with Next-Generation Sequencing experiments: the EMBL Data Science Center, especially Sarah Kaspar and Francesco Tabaro, for support with statistics and bioinformatics, as well as members of the EMBL Rome Epigenetics community for insightful discussions. We thank Ollie Rando and Jamie Hackett for their critical input on the manuscript. Work in this study was supported by EMBL internal funding granted to A.B.

## Author Contributions

Credit attribution. F.S.: Investigation (lead), Formal analysis (lead), Visualization, Writing – original draft. M.R: Investigation. L.C. Investigation. B.D. Investigation, Writing-original draft. A.B: Conceptualization (lead), Investigation, Funding acquisition, Writing – review & editing, Supervision (lead).

## Data Availability Statement

The RNA-sequencing data generated in this study have been submitted to the European Nucleotide Archive (Array Express accession number: E-MTAB-17791) and will be made publicly accessible upon publication. The final accession number will be provided upon acceptance. Data are available to reviewers upon request in the interim. The raw lipidomic data associated with this study is deposited to EMBL-EBI MetaboLights repository under the accession code MTBLS16032, and will be made available upon manuscript acceptance. The code to reproduce the figures from the RNA-seq data presented in this paper is available at https://git.embl.org/grp-boskovic/santilli-et-al-2026-intergenerational-lpd-inheritance.

## Methods

### Animal Husbandry

Animals (*Mus musculus* FVB/NCrl strain) were housed in the Conventional Area within the Laboratory Animal Resources facility on the EMBL Rome “A. Buzzati-Traverso” campus, with separate animal and service areas and strict air-lock entry. Mice were housed under a 12 h light/dark cycle (light from 07:00 to 19:00), with *ad libitum* access to food and water. All experiments involving mice were carried out in accordance with the approved protocols and guidelines by the laboratory animal management and ethics committee of the European Molecular Biology Laboratory (EMBL) under license no. 21-007_RM_AB and Italian Health Ministry license no. 622/2023-PR and 1058/2024-PR.

For the two-generation Low Protein Diet paradigm, experimental males were raised for two generations on CTRL diet, before being assigned to either CTRL or LP diet at weaning (postnatal day 21). All females were raised on CTRL diet. Body weight was measured every three weeks. After 7-9 weeks of continuous dietary intervention, males were paired with females for a maximum of two consecutive days, to minimize their exposure to CTRL diet. If no vaginal plug was detected during the mating period, males were returned to their respective diets for 5 days before reattempting mating with the same female. Upon detection of a vaginal plug, males were euthanized by cervical dislocation, and the entire liver was collected for weight measurement. A portion of the median lobe was subsequently flash-frozen in liquid nitrogen for downstream analyses.

Females were maintained on CTRL diet throughout pregnancy and lactation. The resulting offspring were weighed every three days from birth until weaning, when they were sacrificed by cervical dislocation for liver collection following the same procedures applied to the F0 males.

For the one-generation LP paradigm, F0 animals originated from parents initially weaned on a CHOW diet and later transitioned to CTRL diet. All subsequent procedures were carried out as described for the two-generation paradigm, with the dietary exposure period lasting 7-12 weeks.

### Dietary intervention

Sterilized and tested rodent diets were purchased from Mucedola S.r.l. Briefly, CTRL food is composed of 16.6% of protein and 68.4% of carbohydrates, of which 53% starch and 10% sucrose. The isocaloric LP diet contains 9.2% of proteins with the remaining mass compensated by sucrose, bringing the total carbohydrate content to 75.8%, and the sucrose content to 19%. Consequently, LP food provides fewer protein-derived calories and a higher proportion of energy from carbohydrates. The high cholesterol diet (HCHOL) follows the same composition of CTRL diet, except for the substitution of 2% starch with cholesterol, bringing the total carbohydrate to 66.4%, and making this diet slightly less caloric in terms of metabolized energy.

The full diet composition, and the detailed information on the relative caloric contribution of macronutrients in diet is provided in Supplemental Table 1.

### Weight Measurements and Analyses

F0 males were weighed every three weeks from weaning (3 weeks of age) to sexual maturity (9 weeks) by trained staff of the EMBL Rome LAR. In addition, F1 animals subject to dietary challenge underwent the same weighing protocol, but with weekly measurements. Body weight of F1 offspring was measured every three days from postnatal day 2 to day 20. Downstream analyses were performed using R (v.4.2.1). Statistical comparisons of body weights between experimental groups were conducted by fitting a linear mixed-effects model using the lmer function from the lme4 (v.1.1-35.1) R package. The model included experimental Condition, Age, and their interaction as fixed effects, along with Litter size as a covariate. Cage identity was included as a random effect to account for non-independence of individuals within the same cage post-hoc analysis on the Condition × Age interaction term was used to assess differences between dietary groups at individual time points or across the exposure period. Briefly, estimated marginal means (EMMs) were extracted using the ‘emmeans’ package (v.1.10.0) to perform pairwise comparisons between the experimental groups. To protect against inflation of the Type I error rate due to multiple comparisons, p-values were adjusted using Tukey’s Honestly Significant Difference (HSD) method.

For liver weight analysis, the full liver was weighed, and the liver weight/body weight ratios differences between groups were tested by Wilcoxon rank-sum test (Mann–Whitney–U test), implemented via the stat_compare_means function from the ggpubr (v.0.6.0) R package.

### Liver Transcriptomics

#### Bulk RNA-seq Library preparation

A portion of the median lobe of the liver was dissected from adult F0/F1 (10-15 weeks old, depending on experiment) or young F1 (3 weeks old) animals, and homogenized in Trizol (ThermoFisher, 15596026) using BeadBug™ 6 microtube homogenizer with 1 mm zirconia/silica beads (Roth, N035.1). RNA was extracted using the manufacturer’s recommendations, and subjected to TURBO DNase™ (ThermoFisher, AM2238) treatment according to manufacturer’s recommendations. RNA purity and concentration was quantified using Nanodrop One^C^ (ThermoFisher).

Library preparation for RNA-seq was performed in house, except for the F0 1-gen liver dataset. Briefly, 1 ∝g of starting material was used as input for removal of ribosomal RNA with NEBNext® rRNA Depletion Kit v2 (NEB, E7400). RNA-seq libraries were built using the NEBNext® Ultra™ II Directional RNA Library Prep Kit (NEB, E7760). Quality of libraries was checked with High Sensitivity D1000 ScreenTape assay (Agilent, 5067-5584) on 4150 TapeStation System (Agilent, G2992AA), while concentration was measured using Qubit™dsDNA Quantification Assay Kits (ThermoFisher, Q32854).

Sequencing of the libraries was done by the EMBL Genomics Core Facility, with a 50 bp, pair-end reading mode on an Illumina NextSeq2000 sequencer.

### RNA-seq data analysis

#### Reads processing and Mapping

Raw reads were processed using Trim Galore (v0.6.7) to remove adapter sequences and short, low-quality reads (--quality 23, --length 20, --paired, --stringency 4, and --max_n 1), unless otherwise specified. Reads that passed the quality trimming were aligned to the mouse genome (Genome assembly GRCm38.p6) using STAR (v2.7.11b) in paired-end and reverse-stranded, keeping default mapping parameters. Read quantification was performed directly by STAR using the --quantMode GeneCounts option, which assigns uniquely mapped reads overlapping with exonic regions.

#### Principal component analysis (PCA)

All downstream analysis was performed using R (v4.4.1).

For PCA, read counts after filtering out lowly expressed genes, were transformed using the “vst” function of DESeq2 package (v1.46.0)^37^ in order to make the data homoskedastic (having constant variance along the range of mean values). Transformed counts were input in the “prcomp” function from stats package (v4.4.1).

#### Differential expression analysis

Read counts matrix was used as input for differential expression analysis using DESeq2 (v1.46.0). Variables shared between experimental conditions were included in the design. Genes showing an adjusted p-value < 0.05 and absolute log2 fold change > 0.5 were defined as differentially expressed genes (DEGs) across all comparisons.

#### Over Representation and Gene Set Enrichment Analyses

Differentially expressed genes and gene sets of interest were subjected to over-representation analysis using the “compareCluster” function from the clusterProfiler package (v4.14.6)^57^. To identify enriched terms and pathways, the function was run with the “enrichGO”, “enrichKEGG”, “enrichWP”, and “enrichPathway” options to query the Gene Ontology (GO), Kyoto Encyclopedia of Genes and Genomes (KEGG), WikiPathways (WP), and Reactome databases, respectively. Terms and pathways with adjusted p-value <0.05 were considered statistically significant.

For GSEA genes were ranked by log2 Fold change and input in the function “gseGO”, “gseWP”, “gseKEGG” of clusterProfiler (v4.14.6) on GO, WP, and KEGG databases. Most significant terms for both analyses were visualized using the “dotplot” or “cnetplot” function of enrichplot package (v1.26.6).

#### Gene Overlap Analysis

Venn diagrams were generated using “euler” function from eulerr package (v7.0.4), restricted to genes detected in both datasets. For consistency with our significance threshold, the overlap analysis with the C57BL/6J dataset ^29^ was restricted to DEGs with an absolute log₂ fold change > 0.5. Statistical significance was assessed by one-sided Fisher’s exact test, with p-values adjusted for multiple comparisons using the Benjamini-Hochberg method. Upset plots were generated with the “Upset” function with intersect mode from ComplexHeatmap package (v2.22.0).

#### Gene Expression Correlation Analysis

Spearman rank correlations were computed between LP over CTRL effects (Log2FC) of different experimental groups. Each pairwise correlation was computed using only genes with available data in both groups. Statistical significance was assessed using “cor.test” from stats package (v4.4.1), and 95% confidence intervals (CI) were estimated by percentile bootstrap (1000 resamples), using “boot” and “boot.ci” functions from boot package (v1.3.30).

### Untargeted liver lipidomics

#### Sample preparation

Untargeted lipidomics was performed by the EMBL Metabolomics core facility. Frozen liver samples were homogenized using a bead beater to ensure efficient cell disruption and lipid release. Biphasic extraction of lipids was performed as follows: 80% MeOH (containing internal standards) were initially added to yield a uniform concentration of 50 mg/mL per sample. After homogenization on dry ice via a bead beater (FastPrep-24; MP Biomedicals, CA, USA) at 6.0 m/s (5 x 30 s, 5 min pause time) using 1.0 mm and 2.3mm zirconia/glass beads (Biospec Products, OK, USA), 200 µL of the methanolic suspension were transferred to fresh 1.5 mL laboratory tubes. Subsequently, 500 µL MTBE were added, the monophasic mixture was vortexed for 60 s and incubated at -20 °C for 20 min. For phase separation, 125 µL water was added, followed by another vortexing and incubation step (see previous conditions). The biphasic solvent system was then centrifuged for 10 min at 15,000g and 4 °C with a 5415R microcentrifuge (Eppendorf, Hamburg, Germany). The upper organic phase was transferred, dried under a stream of nitrogen, and reconstituted in 75 µL 90% IPA. Ultimately, the samples were shaken for 10 min, centrifuged (see previous conditions) and supernatants were transferred to an analytical glass vial. The LC-MS/MS analysis was initiated within one hour after the completion of the sample preparation.

Lipidomics and free fatty acids analysis were performed using the ultra-high performance liquid chromatography (Vanquish UHPLC) coupled with Orbitrap Exploris™ 240 high-resolution mass spectrometer (ThermoFisher™) in negative and positive ESI (electrospray ionization) mode.

Chromatographic separation was carried out on an ACQUITY Premier CSH C18 column (Waters; 2.1 mm x 100 mm, 1.7 µm) at a flow rate of 0.3 mL/min. The mobile phase consisted of water:ACN (40:60, v/v; mobile phase phase A) and IPA:ACN (9:1, v/v; mobile phase B), which were modified with a total buffer concentration of 10 mM ammonium acetate + 0.1 % acetic acid (negative mode) and 10 mM ammonium formate + 0.1% formic acid (positive mode), respectively. The following gradient (23 min total run time including re-equilibration) was applied (min/%B): 0/15, 2.5/30, 3.2/48, 15/82, 17.5/99, 19.5/99, 20/15, 23/15. Column temperature was maintained at 65°C, the autosampler was set to 4°C and sample injection volume was set to 1 µL (positive mode) and 3 µL (negative mode). Analytes were recorded via a full scan with a mass resolving power of 120,000 over a mass range from 200 – 1700 m/z (scan time: 100 ms, RF lens: 70%). To obtain MS/MS fragment spectra, data-dependant acquisition was carried out (resolving power: 15,000; scan time: 54 ms; stepped collision energies [%]: 25/35/50; cycle time: 600 ms). Ion source parameters were set to the following values: spray voltage: 3250 V / -3000 V, sheath gas: 45 psi, auxiliary gas: 15 psi, sweep gas: 0 psi, ion transfer tube temperature: 300°C, vaporizer temperature: 275°C.

#### Lipid identification and data analysis

All experimental samples were measured in a randomized manner. Pooled quality control (QC) samples were prepared by mixing equal aliquots from each processed sample. Multiple QCs were injected at the beginning of the analysis in order to equilibrate the analytical system. A QC sample was analyzed after every 5th experimental sample to monitor instrument performance throughout the sequence. For determination of background signals and subsequent background filtering, an additional processed blank sample was recorded.

Lipid identification was performed by matching accurate mass, isotope pattern, MS/MS fragmentation, and retention time using the public MS-DIAL LipidBlast library. Lipid annotations follow the LIPID MAPS nomenclature^58^ to ensure standardized and biologically consistent classification of lipid species. The full list of lipid abbreviations is available as Supplemental Figure 4.

Data was processed using MS DIAL 4.9 and raw peak intensity data was normalized via total ion count of all identified analytes (mTIC). High-confidence (Level 1) feature identification was based on the MS-DIAL LipidBlast V68 library (lipidomics) using accurate mass, isotope pattern, MS/MS fragmentation, and retention time information and a minimum matching score of 90%. To remove misannotations and to enhance confidence in lipid identification, intra-class elution patterns of lipid species were checked for consistency by relying on the expected chromatographic behavior on reversed phase columns within homologous lipid series, considering carbon chain length and degree of saturation.

Only findings that showed a CV <30% in QC samples were considered for further analysis. All downstream processing of the data was performed using R (v.4.4.1).

Statistical analysis was done using limma (v.3.62.2)^59^ by fitting lipid or ontology-wise linear models to log2-transformed, mTIC-normalized intensities, while accounting for relevant experimental covariates. Empirical Bayes procedure was then applied to stabilize the variance estimates across lipids, and differential abundance was assessed using the moderated t-tests. Multiple testing correction to control for type I (false positive) errors was performed using the False Discovery Rate (FDR) method. Lipids with adjusted p-value < 0.05 and absolute log2 fold change > 0.5 were called as differentially abundant.

Lipid Set Enrichment Analysis (LSEA) was performed using a custom workflow inspired by the LipidSigR pipeline^60^ to identify significant structural, functional, and physicochemical trends among altered lipids. First, lipid identifications were converted to shorthand notation and structurally validated using the “parseLipidNames” function of *rgoslin* R package (v.1.10.0). Lipids that could not be parsed were excluded from downstream analyses. Extended lipid characteristics including lipid identity and structure, biophysical and chemical properties, and biological context and functions, were added using the “lipid_annotation” function from *LipidSigR* package (v.1.0.4). To resolve duplicate lipid profiles, only the instance with the highest average log2 intensity was retained for each unique species. Unique lipids were then grouped by shared traits into a custom lipid set library, filtering out terms with fewer than 3 or more than 500 lipids. Finally, this library and the ranked list of lipids by their log2 Fold change, were input to perform enrichment analysis using the “fgseaMultilevel” function of *fgsea* package (v.1.32.4). Statistical significance was evaluated using a Benjamini-Hochberg adjusted p-value threshold of less than 0.05. Enriched terms were categorized into broader biological and chemical groupings for comprehensive visualization via custom bar plots, dot plots, and ridge plots, employing *ggplot2* R package (v.4.0.3).

## Figure Legends

**Supplemental Figure. 1. A.** Boxplot of liver to body weight ratio of males consuming CTRL or LP, at weaning. The p-values were calculated using a two-tailed Wilcoxon rank-sum test (Mann–Whitney U test). **B.** PCA plot of PC1 and PC2 computed on the top 500 variable genes with samples color-coded by litter identity. **C.** Line plots showing the coefficient of determination (adjusted R^2^) from linear regression models quantifying the proportion of variance in each PC explained by individual metadata variables. **D.** PCA plot of PC3 and PC4 computed on the top 500 variable genes, with samples color-coded by dietary group and shaped by litter identity. **E.** Heatmap of Log2 Fold change of the DEGs associated with the significant KEGG pathways identified by over-representation analysis. **F.** Barplot of normalized enrichment score of top 20 significant categories identified by Lipid Set Enrichment Analysis (LSEA). **G.** Boxplot of normalized peak intensities of cholesteryl esters (top), very long acylcarnitinces (middle) and oxidized triglycerides (bottom) in CTRL and LP livers.

**Supplemental Figure. 2. A.** Barplot of the number of animals, grouped by sex and paternal diet. Percentages of males and females withing each experimental group are indicated on top of the bars. **B.** Boxplot of litter sizes grouped by paternal diet. Statistical significance between groups was assessed using a two-tailed Wilcoxon rank-sum test. **C.** Boxplot of liver to body weight ratios of mice sired by CTRL and LP fathers, stratified by sex. Statistical difference between groups was tested by a two-tailed Wilcoxon rank-sum test applied in males and females, separately. Number of animals (n) are indicated at the bottom of the plot. **D.** PCA of PC1 and PC2 computed on the top 500 variable genes with samples color-coded by paternal diet and shaped by biological sex. **E.** Proportion of variance (R^2^) in principal components explained by biological and technical factors. **F.** Expression levels (log2 TPM) of selected genes involved in cholesterol biosynthesis in male and female offspring of CTRL and LP fathers. **G.** Dot plot of significant terms identified by over-representation analysis in the common set of upregulated genes in both F1 LP FVB and C57BL/6J, over the total upregulated DEGs. **H.** Cumulative distribution of relative changes (Log2 Fold change) induced by paternal LP consumption of diglycerides (DG) and triglycerides (TG) compared to the rest of the lipids. **I.** Bar plots of selected ontology-level Log2 fold changes stratified by paternal litter. The number of lipids for each subclass is indicated in brackets.

**Supplemental Figure. 3. A.** PCA based on the 500 most variable genes. Samples are plotted according to their scores on the first two principal components (PC1 and PC2). **B.** Boxplot of normalized expression levels (log2 TPM) of representative urea cycle and cholesterol biosynthesis genes across individual samples grouped by replicate and diet. **C.** Cnetplot showing the connections of the differentially expressed genes enriched in the significant pathways identified by over-representation analysis on Wiki Pathways (left) and KEGG (right) databases. **D.** Venn diagrams showing no overlap between the LP-induced downregulated genes detected in F0 1-gen and F1 FVB/NJ and F1 C57BL/6J. **E.** Heatmap of relative changes induced by LP in Fatty acid (FA) and triglycerides (TG) lipids grouped by number of carbon atoms and number of double bonds. **F.** Heatmap of log2 fold changes for each lipid subclass across biological replicates.

**Supplemental Figure. 4. A.** Growth curves of early postnatal life of animals born from LP or CTRL sires, showing mean body weight (solid line) ± SD (shaded area). T-test on linear mixed model with random and fixed effects was performed for statistical analysis. **B.** Barplot of the number of F1 animals grouped, by sex and paternal diet. **C.** Boxplot of F1 litter size distribution, stratified by paternal diet. Statistical significance between groups was assessed using a two-tailed Wilcoxon rank-sum test. **D.** PCA of top 500 most variable genes from transcriptomic data. **E.** Distribution of transcriptomic sample coordinates along the first six principal components (PC1–PC6), grouped and colored-coded by biological sex (left) and paternal diet (right). **F.** Heatmap of cholesterol biosynthesis genes, showing scaled expression (Z-scores) and relative changes (Log2 Fold change) of LP vs CTRL comparison. **G.** Mean normalized intensity (solid line) ± SD (shaded area) of acylcarnitines grouped by paternal diet and number of carbon atoms. H. Heatmap of log2 fold changes for a subset of lipid ontologies (|log2FC| > 0.5 in at least one replicate) across sex and biological replicates, coloured by LIPID MAPS category.

**Supplemental Figure. 5. A.** Body weight measurements of F1 animals sired by CTRL or LP-fed males (yellow and red lines, respectively) consuming either CTRL, HCHOL or LP diets, stratified by sex. **B.** Representative liver images of F1 animals on three different dietary regimes, indicated, at collection. **C.** Box plots of liver/bodyweight ratios of animals consuming CTRL, LP or HCHOL diets, stratified by paternal diet (CTRL, LP). **D.** Upset plot showing the overlap or unique set of DEGs for F1 dietary effects in paternal CTRL or LP (left), and paternal LP effects in the three different F1 dietary exposures (right) **E.** Volcano plots of all detected genes from liver bulk RNA-seq data of F1 animals sired by LP- or CTRL-fed males, exposed to CTRL (left), HCHOL (middle) or LP (right), after filtering. Genes with adjusted p-values <0.05 & log2 Fold Change > 0.5, were considered significantly differentially expressed (DEGs) in F0 LP vs F0 CTRL testing. **F.** Spearman correlation matrix of the relative effects of paternal LP diet on F1 animals exposed to CTRL, LP, and HCHOL diets.

