## Supplemental Figures S1-S5 for "Cross-generational characterization of paternal Low Protein Diet effects in mice"

### Supplementary Figure S1

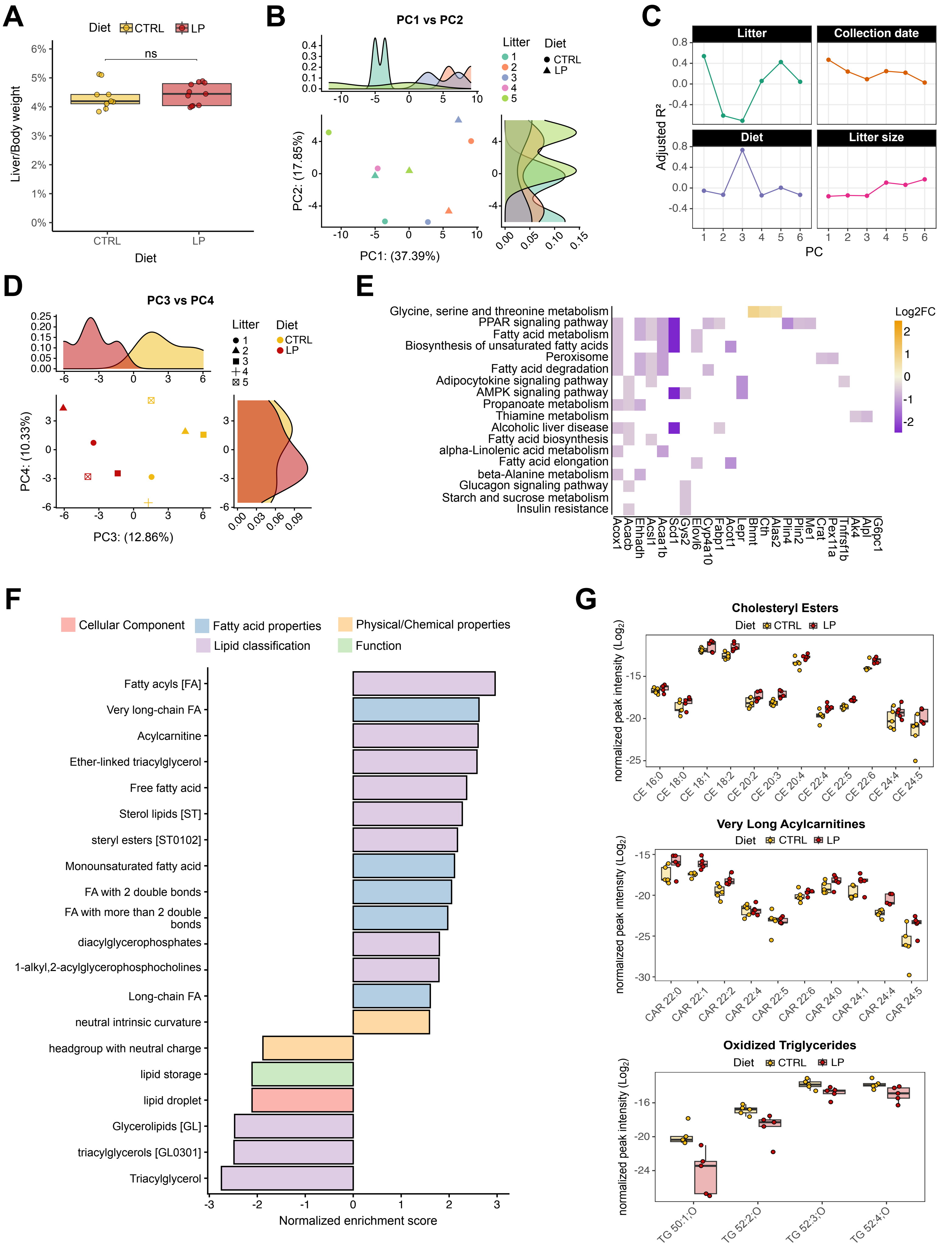

Supplementary Figure S2

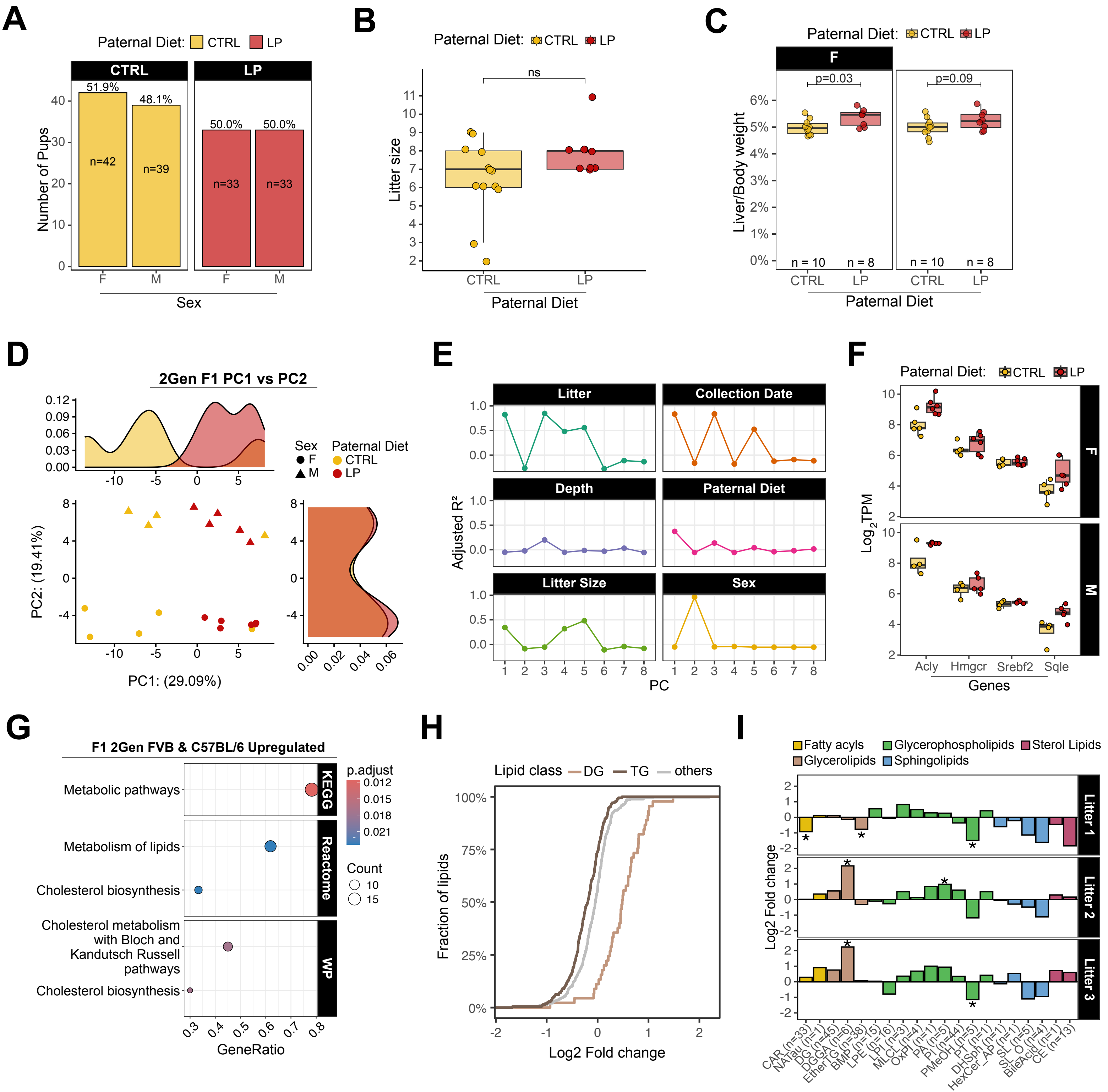

Supplementary Figure S3

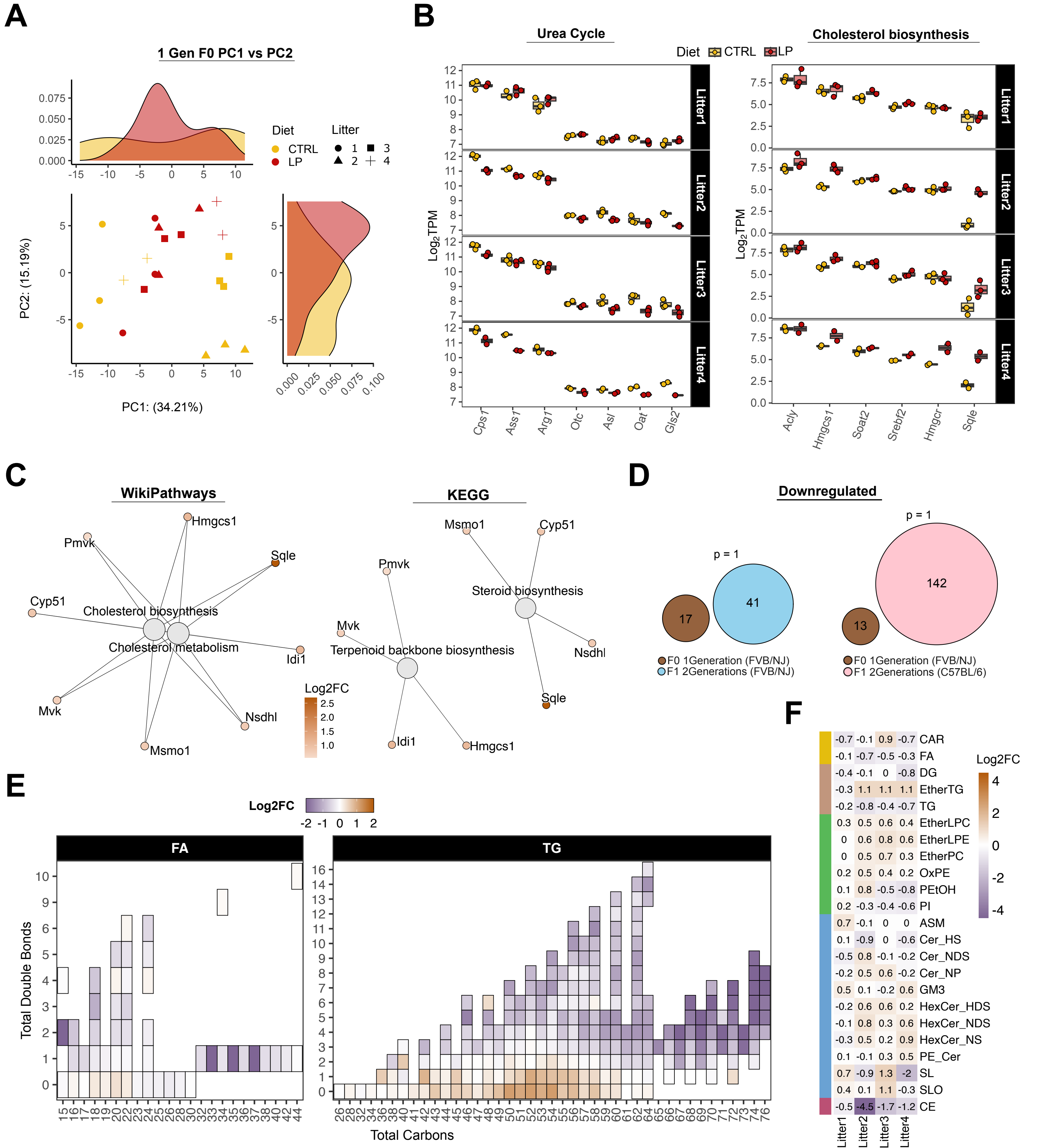

### Supplementary Figure S4

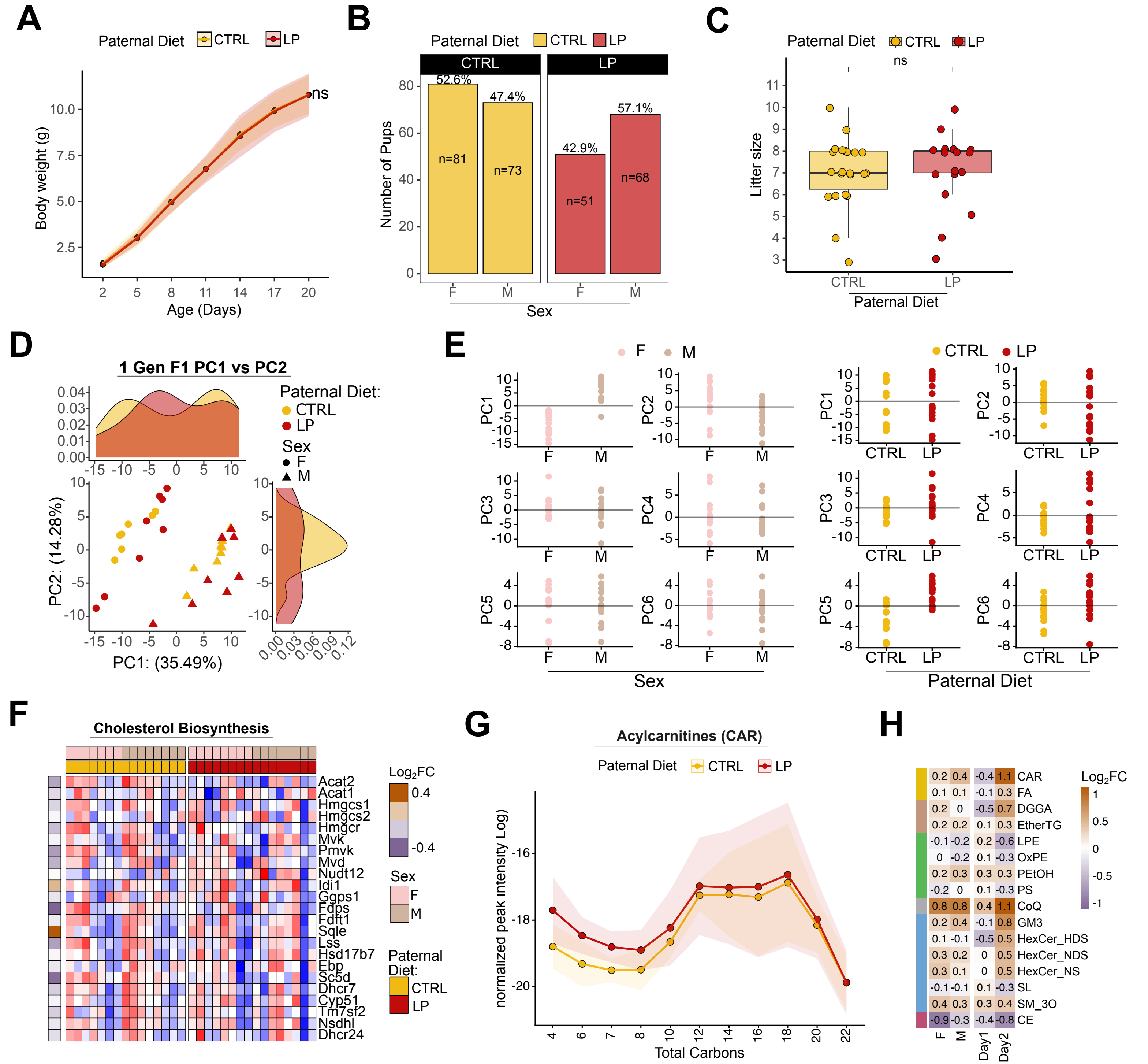

Supplementary Figure S5

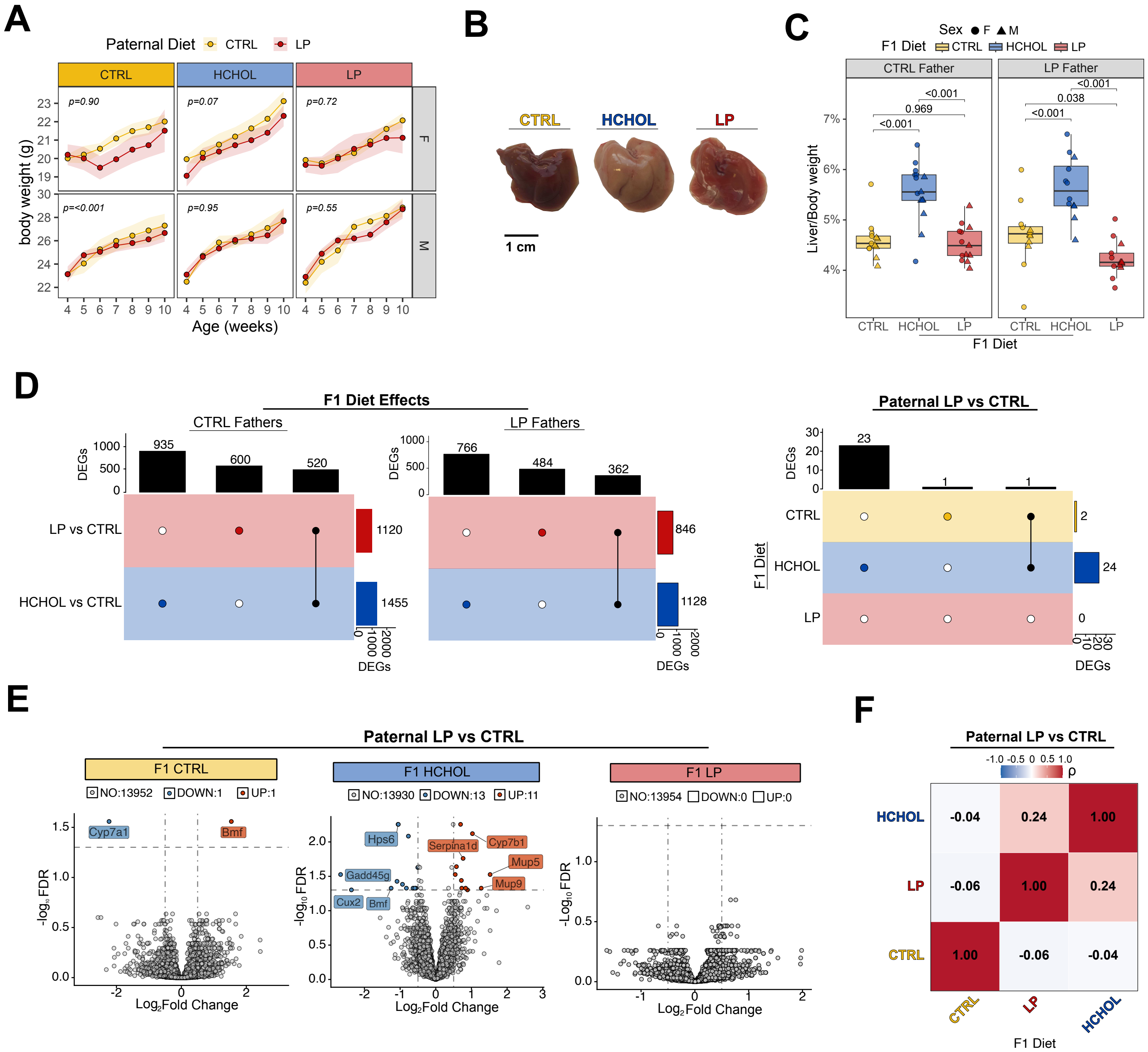
